# Sharing Patterns Between Proteins and Exploring Possible Drug Interactions

**DOI:** 10.1101/2025.10.07.680866

**Authors:** Paloma Tejera-Nevado, Alejandro Rodríguez-González

## Abstract

Due to the nature of amino acids that make up proteins, it is possible to detect small patterns within their sequence. Some of these sequences have been identified as motifs, characterized by defined signatures that are shared across different proteins. In this study, patterns are defined as short amino acid sequences that are commonly found among proteins involved in various types of cancer. The focus of this research is on the localization of such patterns in ALDH2, which is used as a reference protein because it is known to be inhibited by disulfiram. Disulfiram, originally used to treat alcohol use disorder (AUD), has been repurposed for the treatment of lung cancer. Analyzing shared patterns can help identify common protein structures and may assist in locating potential binding cavities and interaction sites. Moreover, it can improve our understanding of how closely related or structurally similar proteins might also be affected by certain drugs. This approach could offer a new perspective on drug specificity and targeting strategies in the context of disease.

## 1 Introduction

### 1.1 Drug-Disease Interactions and Therapeutic Targets

Understanding drug-disease interactions is fundamental to the development of effective and safe therapeutic strategies. These interactions describe how diseases can alter the pharmacokinetics (PK) and pharmacodynamics (PD) of drugs, and conversely, how drugs can influence disease progression or pathology [1]. In complex diseases such as cancer, neurodegenerative disorders, and metabolic syndromes, alterations in cellular signaling pathways, enzyme activity, and gene expression can significantly affect drug responses [2]. This necessitates the identification of therapeutic targets, which are specific molecules or pathways that can be modulated to achieve clinical benefit. Advances in molecular biology, bioinformatics, and systems pharmacology have enabled the discovery of novel targets, such as aberrant enzymes, receptors, and transcription factors, that drive disease mechanisms [3]. Targeted therapies, unlike traditional treatments, offer the potential for precision medicine by minimizing off-target effects and improving efficacy [4]. Analyzing protein-protein interactions (PPIs) is essential for understanding how sequence variations influence disease progression and treatment response. Techniques to study PPIs include experimental and computational methods [5].

### 1.2 Disulfiram as a Multifunctional Therapeutic Agent

Disulfiram inhibits the oxidoreductase enzyme aldehyde dehydrogenase 2 (ALDH2), a crucial enzyme in alcohol metabolism that converts the toxic acetaldehyde into harmless acetate. Inhibition of the ALDH2 by disulfiram leads to acetaldehyde buildup, causing adverse symptoms that deter alcohol consumption, making it effective for treating alcohol dependence [6]. Disulfiram is not specific to ALDH2, as it also inhibits the cytosolic enzyme ALDH1 [7]. Beyond its use in alcohol aversion, disulfiram shows promising anticancer properties, especially in lung cancer [8]. ALDH activity, including ALDH2, is elevated in cancer stem cells (CSC), which contribute to tumor growth, metastasis, and therapy resistance. Disulfiram’s inhibition of ALDH reduces CSC characteristics such as sphere formation, colony growth, and migration, potentially overcoming drug resistance and relapse [9]. Additionally, disulfiram combined with copper induces cancer cell death through apoptosis, ferroptosis, and cuproptosis by generating reactive oxygen species and inhibiting the proteasome system [10]. It can reverse resistance to microtubule inhibitors and has shown antitumor effects in lung cancer models [11]. Clinical trials reveal mixed but encouraging outcomes; a phase IIb trial combining disulfiram with chemotherapy in metastatic non-small cell lung cancer (NSCLC) showed improved progression-free and overall survival, highlighting disulfiram’s role in targeting cancer stem cells [8]. However, further clinical research is needed to confirm its efficacy and best use in cancer therapy [12].

### 1.3 Aldehyde Dehydrogenase (ALDH) family

The human Aldehyde Dehydrogenase (ALDH) family comprises 19 functional genes that encode enzymes responsible for detoxifying aldehydes by oxidizing them into carboxylic acids. These enzymes are distributed across cellular compartments (cytoplasm, mitochondria and endoplasmic reticulum), highlighting their diverse physiological roles. Beyond detoxification, ALDHs are essential in various metabolic pathways, including alcohol metabolism (e.g., ALDH2) and retinoic acid synthesis (e.g., ALDH1A1, ALDH1A2, ALDH1A3), which support cell differentiation and development [13]. ALDH enzymes are significantly involved in disease processes, with disease-causing mutations in the ALDH superfamily falling into three main categories: (1) those that impair NAD binding, (2) those that alter the substrate binding site, and (3) those that disrupt quaternary structure formation [14]. ALDH dysfunction is further implicated in neurodegenerative disorders, including Alzheimer’s and Parkinson’s diseases, due to compromised aldehyde detoxification [15]. Many isoforms are overexpressed in tumors and are linked to cancer stem cell (CSC) traits and resistance to therapy [16]. ALDH1A1, in particular, is a well-established CSC marker in cancers such as lung, breast, and colon, where it contributes to chemotherapy resistance by detoxifying harmful aldehydes [17]. ALDH3A1 is also upregulated in certain cancers, like non-small cell lung carcinoma, and is associated with drug resistance [18].

### 1.4 Protein Sequence Analysis in Drug-Disease Context

Protein sequence analysis is crucial for understanding disease mechanisms and advancing therapeutic development, as proteins are central to cellular function and their dysfunction contributes to many diseases. Aldehyde Dehydrogenase 2 (ALDH2) is a crucial mitochondrial enzyme that functions as a homotetramer. A common variant, ALDH2*2 (Glu504Lys) (present in 30-50 % of East Asian populations) drastically reduces ALDH2 activity and increases cancer risk, especially when combined with alcohol intake [19], [20], [21]. In lung adenocarcinoma (LUAD), ALDH2 downregulation is associated with poorer prognosis, making ALDH2 a potential prognostic factor for early-stage LUAD [22]. Abnormal ALDH activity in cancer is not fully understood but shows promise as a biomarker and therapeutic target. Targeting ALDH, especially in cancer stem cells, could improve treatment outcomes. Repurposed drugs like disulfiram offer potential for safe and effective cancer therapies when combined with conventional treatments [23]. Advances in structure prediction tools like AlphaFold [24] are enabling new methods for protein sequence analysis in drug and disease research. Approaches such as comparing related proteins and identifying conserved patterns in disease-associated sequences [25] aim to reveal shared structural features, including druggable cavities, to support drug repurposing.

This study investigates whether shared amino acid patterns (at least four residues) can reveal functional or structural links between proteins, aiming to distinguish random from meaningful associations and identify off-targets or disease-related candidates. The paper is organized in the following order: Section 2 includes the material and methods used to study protein patterns. In Section 3, the results obtained are presented and discussed. Finally, Section 4 summarizes the conclusions of this study and highlights potential avenues for future research.

## 2 Material and Methods

### 2.1 Classification of Drugs by Action Type, Class, and Gene Symbol

A previous study [25] generated a list of patterns at 5% and 10% occurrence thresholds using a lung cancer treatment dataset, as well as a dataset that included proteins associated with other cancer types: pancreatic (C0346647), breast (C0006142), colon (C0007102), and head and neck (C0010606) cancers. Diseases were identified using their CUI (Concept Unique Identifier), and protein sequence and drug information were obtained from DISNET [26].

First, the data from the lung cancer treatment dataset was analyzed. It was observed that most of the drugs involved are broad-spectrum, which is evident from their action types and associated gene IDs. In total, 43 different drugs were identified. Not all these drugs have a direct correlation with genes specifically related to lung cancer. The drugs were classified according to their action type and the drug class they belong. In this dataset, 12 different action types and 17 drug classes were found. Some drugs have multiple effects and could be valuable for exploring potential interactions with other proteins or targets. However, these multi-effect drugs can also make it difficult to determine whether they are actually binding within a specific protein cavity. A subset of drugs was found to be associated with a single gene symbol, a single action type, and a single drug class. The drug IDs were verified in the ChEMBL database (release ChEMBL_35), and the Drug ID CHEMBL2362016, corresponding to arsenic trioxide, was updated from the original dataset to CHEMBL5483015 in the current update. The molecular structure of the inhibitor disulfiram was retrieved from DrugBank [27].

### 2.2 Analysis and Identification of Shared Sequence Patterns

To investigate the patterns associated with ALHD2 (UniProt entry: P05091) in combination with other proteins, a subset of the data was created focusing on the top five most frequent P05091-containing protein pairs. The cleaned dataset was filtered to extract all the unique sequence patterns associated with each of these protein combinations. For every pair, the number of distinct patterns was computed and stored, along with the list of patterns themselves. The results were then organized into a structured DataFrame to facilitate visualization and interpretation of the co-occurrence pattern data. A search was conducted within the filtered dataset to identify entries containing a specific pattern of interest. When exact matches for the pattern were found, the number of occurrences and corresponding data rows were recorded.

### 2.3 Prediction of Protein Structures and Docking Analysis

Protein sequences were retrieved from UniProt [28]. Protein structure predictions were obtained in PDB format from the AlphaFold DB [29]. However, the structure for protein P30837 (AL1B1) was predicted using the AlphaFold server [30] due to changes in a previous version of the UniProt sequence. Docking predictions were performed using CB-Dock2 [31] and COACH-D [32]. Visualization was carried out with ChimeraX (v 1.6.1) [33], [34], [35]. The Matchmaker tool was used to align the protein structures, and the root mean square deviation (RMSD) was calculated using this software.

## 3 Results and Discussion

### 3.1 Classification of Drugs by Action Type, Class, and Gene Symbol

Focusing on the lung cancer dataset, 43 drugs were detected, many of which are broad-spectrum agents characterized by diverse action types and gene associations. These findings were based on earlier research that identified drug interaction patterns using lung cancer treatment data alongside datasets including proteins from various cancer types (pancreas, breast, colon and head and neck) [25], [36]. The drugs were categorized into 12 action types and 17 drug classes, adding complexity to the analysis. While some drugs exhibit multiple effects that may offer insights into protein interactions, this also complicates pinpointing specific binding sites. Drug entries were filtered to retain only those with a unique drug name, action type, and protein class designation, notably resulting in a subset of drugs that exhibit a one-to-one relationship with a single gene, action type and drug class. This information, derived from the lung cancer treatment dataset, is detailed in Table 1, which lists each drug alongside its corresponding action type, drug class, and gene symbol.

**Table 1.** Filtered Lung Cancer Treatment Dataset.

| Drug id | Drug name | Action type | Protein Class name | Gene symbol |
| --- | --- | --- | --- | --- |
| CHEMBL185 | Fluorouracil | Inhibitor | Transferase | TYMS |
| CHEMBL5483015 | Arsenic trioxide | Inhibitor | Oxidoreductase | TXNRD1 |
| CHEMBL238071 | Vindesine | Inhibitor | Cytoskeletal protein | TUBB1 |
| CHEMBL56367 | Nimesulide | Inhibitor | Oxidoreductase | PTGS2 |
| CHEMBL924 | Zoledronic acid | Inhibitor | Transferase | FDPS |
| CHEMBL964 | Disulfiram | Inhibitor | Oxidoreductase | ALDH2 |

This filtering strategy was applied to ensure specificity and reduce the number of candidates for validation. From this approach, six drugs met the criteria. ALDH2 was selected as a primary candidate due to its known history in drug repurposing. For example, disulfiram (CHEMBL964), a drug originally developed for alcohol use disorder, has been investigated for the treatment of non-small cell lung cancer. Disulfiram works by inhibiting ALDH2. Additionally, arsenic trioxide and nimesulide were identified as inhibitors targeting the oxidoreductase class proteins TXNRD1 and PTGS2, respectively.

### 3.2 Analysis of Sequence Patterns in ALDH2

The analysis began by identifying the number of sequence patterns shared between ALDH2 (Uniprot entry: P05091) and other proteins. In total, 90 patterns were detected, with one pattern (“AKLL”) falling under the 10% occurrence threshold and the remaining 89 corresponding to the 5% occurrence category. Subsequently, patterns not found in other cancer types were excluded from the dataset. This filtering step led to the removal of only two patterns, resulting in a final set of 88 patterns for further analysis in the case of ALDH2. To further explore potential co-associations involving ALDH2, the top five protein combinations containing P05091 were selected for analysis. For each combination, the dataset was filtered to identify all unique patterns associated with the pair. The total number of distinct patterns per combination was then calculated. To define a reference framework for pattern comparison, proteins sharing at least two patterns with ALDH2 were identified, yielding five protein candidates. The results, including the number of pattern occurrences and the specific patterns identified for each protein pair, are summarized in Table 2.

**Table 2.** Sequence Patterns Associated with ALDH2 (P05091) Paired with Selected Proteins.

| Protein id | Gene symbol | Occurrences | Patterns |
| --- | --- | --- | --- |
| P42345 | MTOR | 9 | [AALET, AAFQ, ANYL, ELGE, FQLG, KTEQ, QIFI, QQPE, VLKC] |
| P22102 | GART | 6 | [AWKL, EDVD, GQII, LDKA, VARA, VDLD] |
| Q16881 | TXNRD1 | 6 | [GLAAA, GDKE, LQAG, LQAY, RVVG, TEVK] |
| P51649 | ALDH5A1 | 5 | [FTGST, PWNFP, DAVS, PFGG, VAEQ] |
| P10696 | ALPG | 4 | [AAARF, FGGY, KLGP, SDAD] |

TXNRD1 was prioritized for initial inspection because it was also present in the lung cancer treatment dataset. The protein TXNRD1 shares six patterns with ALDH2, based on data from the treatment dataset, which was used in this case to assess the effects reported in the references. A manual inspection of the shared patterns was not located within the ligand-binding cavity of ALDH2, nor were they associated with structurally similar regions between ALDH2 and TXNRD1 based on predicted protein folding models. This suggests that while sequence patterns can indicate potential biological relationships, not all shared patterns correspond to functional or structural similarity in drug interaction sites. A similar observation was made when examining the common patterns detected between the pairs ALDH2 and MTOR, GART, and ALPG (data not shown).

As part of efforts to identify therapeutic targets in cancer, analysis of the lung cancer treatment dataset used in this study revealed that both ALDH2 and ALDH5A1 are implicated in lung cancer therapy. Disulfiram, an FDA-approved drug originally used for alcohol aversion therapy, acts as an inhibitor of ALDH2 and is under investigation for its potential in treating non-small cell lung cancer [10]. Valproic acid, on the other hand, inhibits ALDH5A1 [37]. Disulfiram is also gaining considerable attention for its broader anticancer potential, largely due to its ability to inhibit aldehyde dehydrogenase 1 family member A1 (ALDH1A1) [38]. ALDH1A1 is often overexpressed in various cancers and is linked to tumor aggressiveness, drug resistance, and the survival of cancer stem cells (CSC) [39], [40]. To explore potential functional similarities, shared amino acid patterns were analyzed between two mitochondrial enzymes, ALDH2 (mitochondrial aldehyde dehydrogenase, 517 amino acids) and ALDH5A1 (mitochondrial succinate semialdehyde dehydrogenase, 535 amino acids). The analysis revealed that the sequences “FTGST” and “PWNFP” are located within regions predicted to form part of the ligand-binding cavity, suggesting possible overlap in substrate interaction or inhibitor binding. Additionally, the pattern “VAEQ” was identified in both proteins within two distinct regions also associated with the ligand-binding site, while the patterns “DAVS” and “PFGG” were found outside the surrounding area of the ligandbinding site (**¡Error! No se encuentra el origen de la referencia**.).

Disulfiram irreversibly inactivates ALDH enzymes by carbamylating a key cysteine residue in their active site [38], [41]. While disulfiram broadly inhibits ALDHs, including ALDH2 (relevant to its anti-alcohol effects), its action on ALDH1A1 specifically targets a pathway vital for cancer progression [42]. Studies demonstrate that disulfiram, particularly when combined with copper, effectively reduces CSC populations, inhibits tumor growth, and enhances the sensitivity of cancer cells to chemotherapy across various cancer types, including lung, ovarian, and glioblastoma [39], [43], [44]. Beyond ALDH inhibition, disulfiram’s broader therapeutic effects in cancer are being explored, encompassing its ability to induce various forms of regulated cell death, such as apoptosis, ferroptosis, and cuproptosis, and to modulate other critical cellular signaling pathways [45], [46], [47].

### 3.3 Proteins Sharing the Selected Sequence Pattern

Focusing on proteins involved in cancer and drug interaction, the aim was to identify conserved amino acid patterns that may indicate shared structural or functional features. The identification of shared residues was guided by reference data from protein treatments, aiming to reveal potential common protein structures and binding cavities. In this case, five of the common patterns detected were found to belong to ALDH2 (P05091) and ALDH5A1 (P51649), with the latter corresponding to the protein SSDH. A search was conducted to identify proteins containing the specific pattern “FTGST” within the dataset comprising proteins from other cancer types, such as pancreatic (C0236647), breast (C0006142), colon (C0007102), and head and neck (C0010606) (Table 3).

**Table 3.** Proteins from Other Cancer Types Containing the Pattern “FTGST”.

| Protein id | Protein name | Disease id | Proteins Treat. id | Protein name Treat. |
| --- | --- | --- | --- | --- |
| P00352 | AL1A1 | C0006142 | P05091 | ALDH2 |
|  |  | C0346647 | P51649 | SSDH |
| P47895 | AL1A3 | C0006142 |  |  |
| Q9Y6D5 | BIG2 | C0006142 |  |  |
| Q9Y6D6 | BIG1 | C0006142 |  |  |
| P30837 | AL1B1 | C0007102 |  |  |
| Q14031 | CO4A6 | C0007102 |  |  |
| P18583 | SON | C0010606 |  |  |

An initial inspection of the dataset identified AL1A1, AL1A3, and AL 1B1 as proteins closely related to ALDH2, and they were analyzed further. The proteins BIG2 and BIG1 were also identified, corresponding to Brefeldin A-inhibited guanine nucleotide-exchange protein 2 (1785 aa) and type 1 (1849 aa), respectively. These two proteins share a similar folding structure; however, this was not related to ALDH2 (data not shown). The proteins CO4A6 and SON correspond to Collagen alpha-6(IV) chain (1691 aa) and protein SON (2426 aa), respectively. Due to their notably unstructured regions, they were not evaluated further (data not shown).

The next pattern selected for detailed analysis was “PWNFP”, chosen to explore further its occurrence and potential functional relevance among the proteins detected in other cancer types (Table 4). The pattern “PWNFP” is detected only in the related proteins AL1A1, AL1A3 and AL1B1. The protein AL1A1 is associated with breast (C0006142) and pancreatic (C0346647) cancer. The protein AL1A3 is linked to breast cancer (C0006142), while AL1B1 is related to colon cancer (C0007102).

**Table 4.** Proteins from Other Cancer Types Containing the Pattern “PWNFP”.

| Protein id | Protein name | Disease id | Proteins Treat. id | Protein name Treat. |
| --- | --- | --- | --- | --- |
| P00352 | AL1A1 | C0006142 | P05091 | ALDH2 |
|  |  | C0346647 | P51649 | SSDH |
| P47895 | AL1A3 | C0006142 |  |  |
| P30837 | AL1B1 | C0007102 |  |  |

Subsequently, ALDH2, which belongs to the treatment dataset and is associated with disulfiram, was used as a reference. AL1A1, AL1A3 and AL1B1 were the proteins selected to localize the pattern “FTGST” among all the detected proteins (Fig. 2A). Additionally, the pattern “PWNFP” was detected in these proteins, as they were the ones in which this pattern appeared (Fig. 2B). Structural localization of both “FTGST” and “PWNFP” in these proteins, using ALDH2 as a reference, confirmed their presence in comparable regions (Fig. 2A, 2B). These findings support the hypothesis that shared patterns across ALDH family members may relate to conserved functions and potential drug-binding sites. Current research also focuses on developing more selective ALDH1A1 inhibitors derived from disulfiram to maximize therapeutic benefits while minimizing off-target effects [38].

**Fig. 1.**
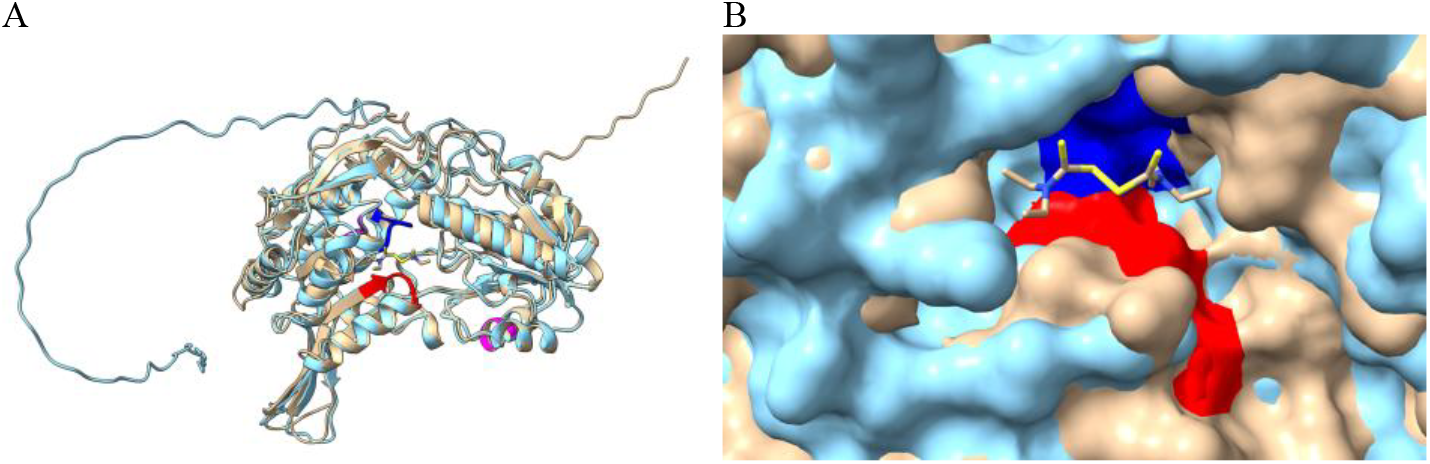
Structural comparison of ALDH2 (UniProt ID: P05091, shown in brown) and ALDH5A1 (UniProt ID: P51649, shown in cyan), highlighting shared sequence patterns. (A) Overall structural overview. (B) Detailed view of the region surrounding the disulfiram-binding site, generated using CB-Dock2. The pattern “FTGST” (colored red) is located at position 260 in ALDH2 and 282 in ALDH5A1, positioned at the end of a β-sheet and extending into an unstructured region; the pattern appears to overlap spatially between the two structures. The pattern “PWNFP” (colored blue), found at position 184 in ALDH2 and 203 in ALDH5A1, is also situated in an unstructured region and shows spatial proximity between the proteins. In contrast, the patterns “DAVS” and “PFGG” are not located near the ligand-binding region and do not show close spatial alignment. The pattern “VAEQ” (colored magenta) appears at position 210 in ALDH2 and 481 in ALDH5A1; although not directly adjacent to the binding cavity, it is located nearby in both structures within unstructured regions. Structural alignment using the Matchmaker tool yielded an RMSD of 0.856 Å over 394 pruned atom pairs, indicating a high degree of structural similarity. The visualization was performed using UCSF ChimeraX.

**Fig. 2.**
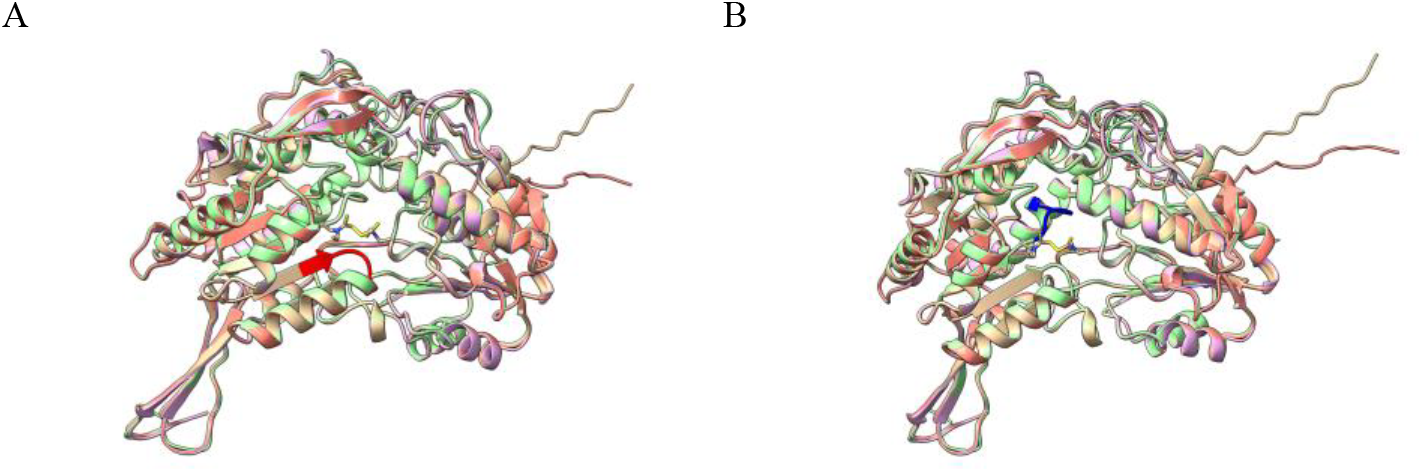
Comparison of predicted protein structures between ALDH2 and proteins sharing common patterns in other cancer types. ALDH2 (brown) was used as a reference, and a structural comparison was performed with ALA1A1 (pink), ALA1A3 (green), and AL1B1 (salmon). The pattern “FTGST” is marked in red (A), and “PWNFP” is colored in blue (B). Visualization and structural overlap were conducted using the MatchMaker tool in ChimeraX.

### 3.4 Analysis of Pattern Localization in Protein Active Sites

The patterns shared by some proteins did not contain ant cysteine residues that could be involved in the inhibition of ALDH2 by disulfiram. Therefore, the study proceeded with a search for the catalytic cysteine at position 302 and adjacent cysteine residues. The pattern “CCAG” was identified, and the proteins containing this pattern were further analyzed (Table 5). In this case, the protein associated with the treatment condition corresponds to ALDH2, and PSMD1 also shares that pattern.

**Table 5.** Proteins from Other Cancer Types Containing the Pattern “CCAG”.

| Protein id | Protein name | Disease id | Proteins Treat. id | Protein name Treat. |
| --- | --- | --- | --- | --- |
| Q13077 | TRAF1 | C0006142 | P05091 | ALDH2 |
|  |  | C0007102 | Q99460 | PSMD1 |
| Q9H2E6 | SEM6A | C0006142 |  |  |
|  |  | C0007102 |  |  |
| Q96GP6 | SREC2 | C0006142 |  |  |
| P30837 | AL1B1 | C0007102 |  |  |
| O75508 | CLD11 | C0007102 |  |  |

The localization of the pattern “CCAG” and the structural visualization of all proteins were performed, with further analysis focusing on the binding cavity of ALDH2 with disulfiram. For the docking analysis, the full sequences of the proteins were used. The predicted complex structure was then determined using COACH-D and compared to that of AL1B1, the protein with the most similar molecular structure. Structural localization showed that “CCAG” lies in an unstructured region of the protein (Fig. 3).

**Fig. 3.**
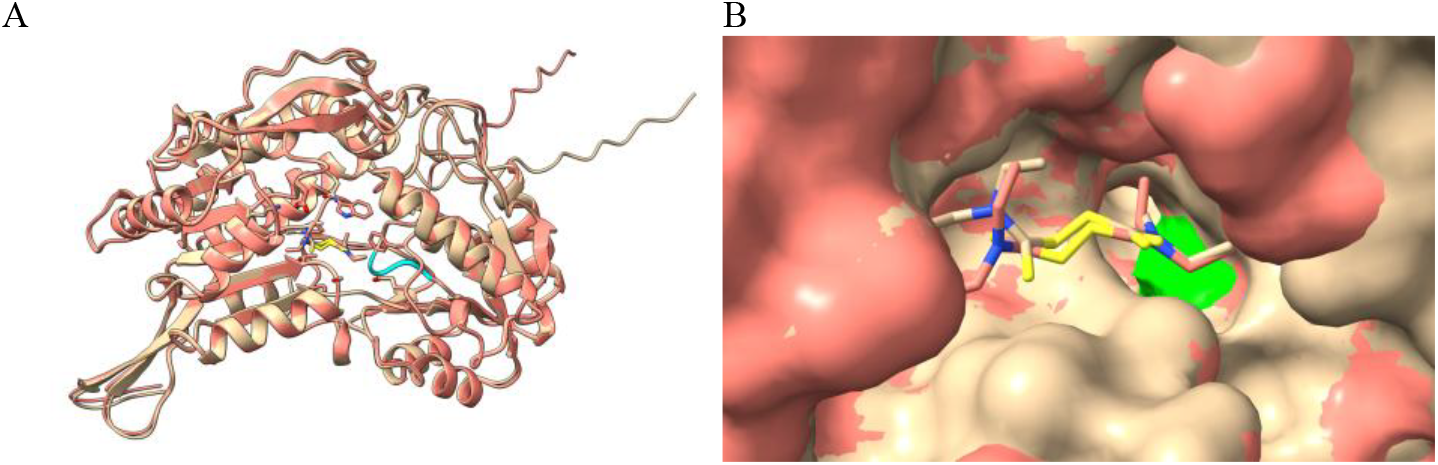
Comparison of predicted protein structures between ALDH2 and AL1B1. ALDH2 (colored in brown) is used as the reference protein for the complex with disulfiram and is compared to AL1B1 (colored in salmon). (A) The pattern “CCGA” is highlighted in cyan. (B) The cysteine at position 309, which is the first “C” in the “CCAG” pattern, is marked in green and is localized within the binding cavity. Visualization was performed using ChimeraX.

It is known that the first 18 amino acids of ALDH2 correspond to the mitochondrial targeting signal, which is cleaved post-translationally. Consequently, the predicted binding residues are shifted by 17 positions. The results reveal that a cysteine residue is identified in both ALDH2 and AL1B1 as part of the predicted binding sites involved in the interaction with disulfiram. Following this, the complex information for ALDH2 and AL1B was analyzed, including the C-Score, TM-score, and the predicted binding sites of the top-ranked model (Table 6).

**Table 6.**
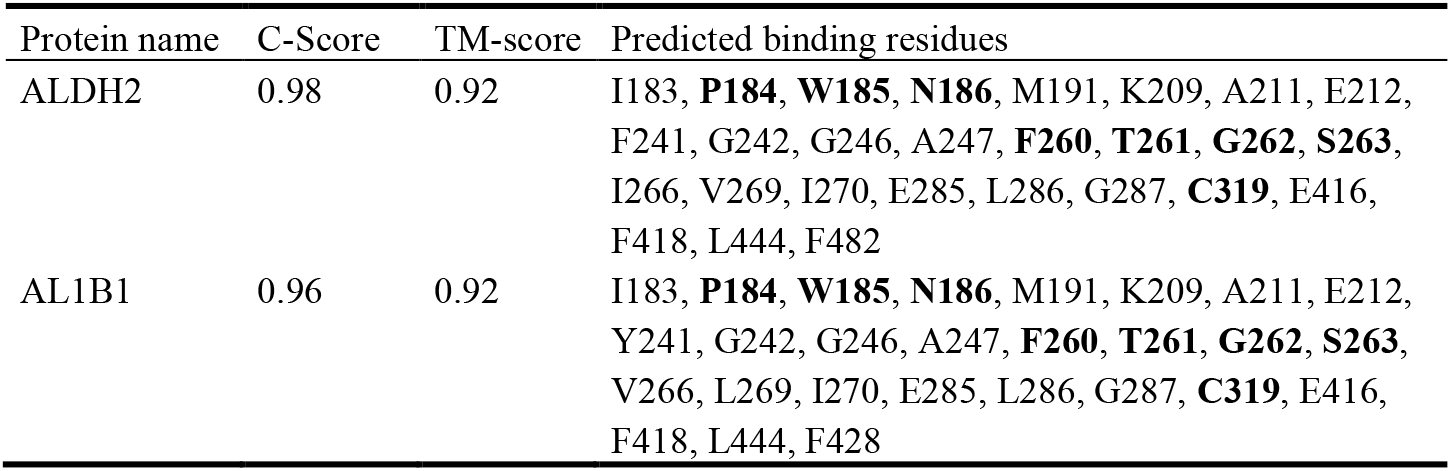
COACH-D Results for ALDH2 and AL1B1 with Disulfiram.

| Protein name | C-Score | TM-score | Predicted binding residues |
| --- | --- | --- | --- |
| ALDH2 | 0.98 | 0.92 | I183, **P184**, **W185**, **N186**, M191, K209, A211, E212, F241, G242, G246, A247, **F260**, **T261**, **G262**, **S263**, I266, V269, I270, E285, L286, G287, **C319**, E416, F418, L444, F482 |
| AL1B1 | 0.96 | 0.92 | I183, **P184**, **W185**, **N186**, M191, K209, A211, E212, Y241, G242, G246, A247, **F260**, **T261**, **G262**, **S263**, V266, L269, I270, E285, L286, G287, **C319**, E416, F418, L444, F428 |

ALDH2 and AL1B1 share 72.7% sequence identity over an overlap of 517 residues, and within the predicted binding sites, the amino acids “PWN” and “FTGS” (part of the patterns “PWNFP” and “FTGST”) are present, along with cysteine 302 (position 319 in the full sequence), which is part of the “CCAG” pattern and included in the predicted disulfiram binding site, highlighting this region’s potential relevance in drug binding. AL1B1 is associated with colon cancer. The combination of disulfiram and copper significantly enhances its anti-tumor efficacy by targeting ALDH, the NK-κb pathway, the MAPK pathway, and other mechanisms [48]. Additionally, the disulfiram/copper complex has been shown to potentiate the anti-tumor effects of 5-fluorouracil in colon cancer models [49].

Aldehyde dehydrogenases (ALDHs) are a family of NAD(P)^+^-dependent enzymes involved in the oxidation of aldehydes to their corresponding carboxylic acids, playing essential roles in cellular detoxification, metabolism, and protection against oxidative stress [50]. Among them, ALDH2 is a key mitochondrial isoform primarily responsible for metabolizing toxic acetaldehyde generated during alcohol metabolism [51]. Structurally, ALDH2 functions as a homotetramer, with each subunit comprising three domains: a catalytic domain, a coenzyme-binding domain, and an oligomerization domain. The active site, located at the interface of subunits, contains the catalytic residue Cys 302, which is critical for enzymatic activity [13]. The tetrameric organization is necessary for structural integrity and optimal catalytic function. Another mitochondrial member of the ALDH family is AL1B1, which shares significant sequence and structural similarity with ALDH2. Like ALDH2, AL1B1 contains a mitochondrial targeting sequence at its N-terminus. This signal peptide directs the protein to the mitochondria and is cleaved post-translationally, resulting in the mature, active form of the enzyme localized within the mitochondrial matrix. The mitochondrial localization of both ALDH2 and AL1B1 underscores their involvement in regulating intracellular aldehyde levels and their potential relevance as therapeutic targets, particularly in diseases where aldehyde accumulation and oxidative stress are pathogenic factors [52]. The therapeutic inhibition of ALDH enzymes (particularly with broad-spectrum agents like disulfiram) offers potential benefits across diseases such as cancer and neurodegeneration [14], [53]. However, ALDH’s essential role in detoxification and cellular stress response means that non-specific or long-term inhibition may inadvertently drive drug resistance by enriching ALDH-overexpressing, therapy-resistant cells, including cancer stem cells. This underscores the need for greater precision in drug-target interactions, favouring isoform-selective inhibitors that minimize off-target effects.

## 4 Conclusion and Future work

In this study, patterns are defined as short amino acid sequences that are shared between different proteins. Some of these patterns are associated with specific protein families, while others may arise due to the probability of certain amino acids appearing together, given 20 standard amino acids found in proteins. The main objective was to evaluate whether these shared patterns could be used to identify new drug repurposing candidates or to predict potential off-target protein interactions. To investigate this, we focused on inhibitors, proteins, and patterns identified in a dataset related to the treatment of lung cancer. Among the proteins analyzed, several belong to the aldehyde dehydrogenase (ALDH) family, which is of particular interest due to its known role in cancer metabolism and drug resistance. These include ALDH2, AL1A1, AL1A2 and AL1B1, which were consistently represented across the datasets. The presence of conserved patterns within this protein family may point to shared structural or functional features that are relevant in the context of cancer progression and therapeutic targeting. The separation of ALDH family proteins in this analysis is crucial, as their conserved patterns may not only reflect evolutionary relationships but also indicate common binding sites or functional domains. This insight can guide future studies aimed at understanding how repurposed drugs, such as disulfiram, interact with members of the ALDH family and potentially influence treatment outcomes across multiple cancer types. This also underscores the potential of such drugs to exhibit broad-spectrum activity by targeting conserved patterns among related proteins, possibly extending their effect beyond a single protein to entire families or interaction networks.

Future work should focus on a deeper characterization of expression patterns of ALDH isoforms and other motif-bearing proteins across different cancer types and cellular contexts. Understanding how these proteins are differentially expressed in specific diseases and cell populations could improve predictions of drug sensitivity and resistance. Integrating expression data with structural and functional pattern analysis may ultimately support the development of more precise, context-specific therapeutic strategies.

## Acknowledgments

The work is a result of the project “Data-driven drug repositioning applying graph neural networks (3DR-GNN)” that is being developed under grant “PID2021-122659OB-I00” from the Spanish Ministry of Science and Innovation.

## Disclosure of Interests

The authors have no competing interests to declare that are relevant to the content of this article.

## Notes

### Competing Interest Statement

The authors have declared no competing interest.

